# Probing learning through the lens of changes in circuit dynamics

**DOI:** 10.1101/2023.09.13.557585

**Authors:** Owen Marschall, Cristina Savin

## Abstract

Despite the success of dynamical systems as accounts of circuit computation and observed behavior, our understanding of how dynamical systems evolve over learning is very limited. Here we develop a computational framework for extracting core dynamical systems features of recurrent circuits across learning and analyze the properties of these meta-dynamics in model analogues of several brain-relevant tasks. Across learning algorithms and tasks we find a stereotyped path to task mastery, which involves the creation of dynamical systems features and their refinement to a stable solution. This learning universality reveals common principles in the organization of recurrent neural networks in service to function and highlights some of the challenges in reverse engineering learning principles from chronic population recordings of neural activity.

## Introduction

It is generally accepted that synaptic plasticity is the primary physiological driver of learning. However, making direct connections between changes in synaptic strength and changes in behavior is difficult, as they occur on vastly different spatial and temporal scales. Circuit dynamics can act as a bridge between the synaptic and behavioral levels of description: synaptic plasticity drives changes in circuit dynamics, and changes in circuit dynamics drive changes in behavior (1). Furthermore, recent methodological advancements such as large-scale chronic population recordings (2) and statistical tools for their analysis (3) allow for unprecedented access to this level. However, we are missing a principled account for how learning at the behavioral level manifests as changes in circuit dynamics.

The past decades have brought significant progress in linking circuit dynamics to function. Experimental and theoretical work have come together to identify an array of dynamical systems features that offer compact computational characterizations of behavior (4, 5). In particular, low-dimensional attractor dynamics are ubiquitous in the brain, across circuits and species (6). They come in several flavors, either single discrete states (fixed points) or multiple states that effectively behave as a continuum (attractor manifolds), and support essential brain functions, from short-term (7) and associative memory (8), to denoising (6), cognitive map formation (9), integration (10), and decision making (11). Attractor dynamics also provide powerful means for interpreting how recurrent neural networks (RNNs) solve complex tasks (12).

While it is arguably difficult to link phenomenological models of synaptic plasticity to behaviorally relevant function in a bottom-up fashion, top-down models of plasticity directly map global behavioral objectives to learning rules. In particular, RNNs can be trained by optimizing a task-specific objective function through either machine learning algorithms (13, 14) or biologically motivated approximations thereof (15). Such trained RNNs provide a useful tool for reasoning about brain computation (16, 17). They are also a fruitful test-bed for investigating biological learning (18, 19).

Here we develop a theoretical framework for assessing learning through the lens of changes in population activity. Our premise is that dynamical systems features such as fixed points or manifold attractors have explanatory power at the level of behavior, and measurable signatures in neural population recordings (7, 20–22) or RNN activity (12, 23). Thus, tracking the evolution of these features over learning, i.e. the network “meta-dynamics,” offers an indirect lens into the learning process—ultimately driven by changes in synaptic connectivity—which links explicitly to observed changes in behavior. The main practical advantage of this formulation is that neural activity can be experimentally observed and manipulated *in vivo* with greater precision than synapses.

Using RNNs as a model system for testing our ideas, we develop tools for identifying the dynamical systems features present at any moment and metrics for quantifying changes in dynamical systems structure over learning. We apply our approach to several tasks that capture key brain computations such as item working memory, evidence integration and decision making. Beyond idiosyncrasies of different tasks and algorithms, our analysis reveals common patterns of meta-dynamics over learning, where qualitative changes in dynamical systems structure accompany breakthroughs in task performance. These results argue that we should re-center questions about biological learning on network dynamics, despite the nominal importance of synaptic changes in the formulation of learning rules.

## Results

We start by training recurrent neural networks to perform a simple, online version of a working memory task, specifically the three dimensional “Flip-Flop” task (Fig 1b, 24). Networks receive 3 sparse, independent input streams 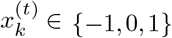. Non-zero inputs need to be maintained persistently until a conflicting input is presented, which “flips” the memory state and the corresponding output 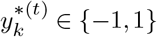. There is a “universal” solution to this task (25) in terms of the internal network dynamics, involving 8 fixed-point attractors organized in a cube with symmetric input-dependent transitions between them (Fig 1b). This solution is robustly shared across architectures, learning dynamics and networks. Our fundamental question is, how does this precise structure emerge through learning?

**Fig. 1.**
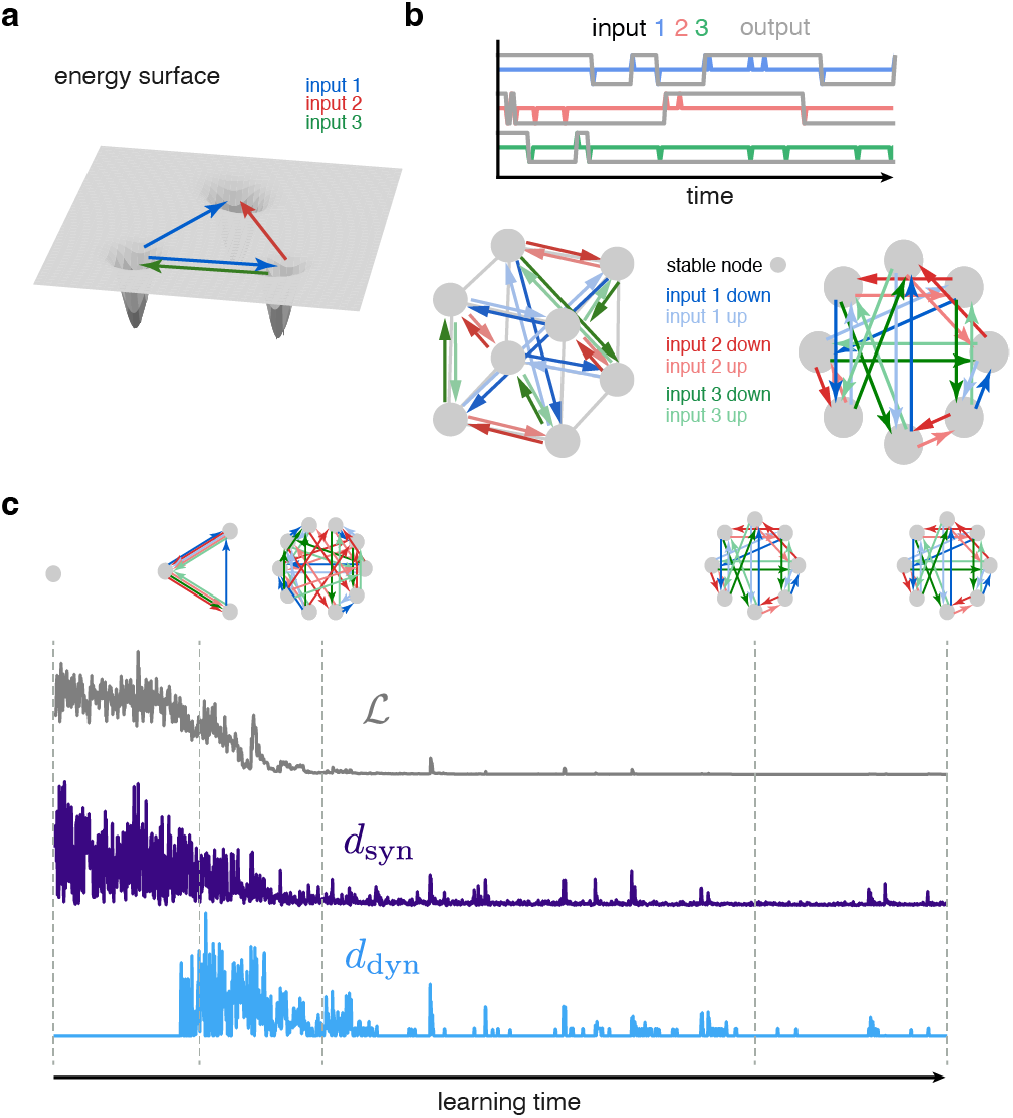
**a)** Cartoon illustrating how the relationships between fixed points in a network’s dynamics can relate to one another through task-relevant inputs. Inputs can drive the network from one fixed point to the basin of attraction for another. **b)** The Flip-Flop task and its universal solution. Each of 3 dimensions is color-coded (blue, red, green), with correct outputs in grey, which each flip in response to the non-zero pulses of its input channel. The left multigraph shows the fixed points and input driven-transitions of the universal solution as embedded in a low-dimensional projection of the network state space. The right multigraph abstracts the version of the left away from the geometric embedding, as a topological entity. **c)** Snapshot of the learning process, via the test loss ℒ (grey), the rate of change in the recurrent weights *d*_syn_ (purple) and the rate of change in topological structure *d*_dyn_. At sam-ple times of t = 0, 3200, 6000, 15660, 19990, we show what the network topology looks like at this point in learning.

### Learning through the lens of meta-dynamics

In our circuit models, learning manifests through changes in the synaptic strengths **W** = [**W**^rec^, **W**^in^], which define the network dynamics via **r**^(*t*)^ = *ϕ*(**W**^rec^**r**^(*t−*1)^ + **W**^in^**x**^(*t*)^). We build on the approach from (12, 25) to precisely reverse-engineer the RNN as a function of learning time. For a given **W**, the fixed points correspond to stable local minima of the network’s kinetic energy 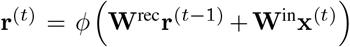, which can be calculated numerically (see Methods). Once fixed points are determined, the input-driven transitions among them are obtained by tracking the evolution of the network dynamics for different initial conditions and inputs. These dynamical properties of a given network can be compactly summarized in a directed *multigraph* structure, *T*_*pqr*_ where nodes *p* and *q* are stable fixed points, and edges of a given type describe transitions for an associated input *r*. This multigraph is unique to our approach. It abstracts away the exact form of the basins of attraction and the geometric alignment of input vectors that allows for transitions between them, and provides a complete characterization of the network’s mechanism for solving the task.

Although changes in synaptic strength ultimately drive these improvements in functionality over learning (via any given plasticity rule), the relation between the two is not straightforward. In particular, there is a period early in training with large overall changes in synaptic strengths, i.e. *d*_syn ≡‖_**W**^(*t*+Δ*t*)^ **W**^(*t*)^ _‖ ≫_0, but little change in overt behavioral performance, as quantified by ℒ(Fig. 1c). Only once new useful underlying dynamical systems features (in this case fixed point attractors) emerge, does the network start improving task performance. Thus it is via changes in the *mechanism*—as summarized by the multigraph **T**—that synaptic plasticity eventually leads to task learning. Therefore the evolution of the dynamics themselves, the *meta-dynamics*, is a more direct avenue to study the process of learning as it pertains to measureable changes in behavior.

Snapshots of the network mechanism over time provide a picture for how the network function evolves over learning (Fig 1c). This can be compactly summarized via a metric that quantifies the *rate* of dynamical change from moment to moment, by how dissimilar the multigraphs **T**^(*t*)^ and **T**^(*t*+Δ*t*)^ are between successive snapshots of the learning process. To quantify the differences between two multigraphs, the *dynamical distance*, requires a principled alignment of their nodes. As the number of nodes changes over learning, it is nontrivial to track the identity of a given fixed point between two multigraphs, but the ambiguity can be largely resolved by a fixed point’s associated behavioral readout (see Methods). Given a correspondence between nodes, the **rate of dynamical change** *d*_dyn_(*t*) is defined as the (negated) dot-product similarity between the input-driven transition patterns **T**^(*t*)^ and **T**^(*t*+Δ*t*)^. These values are shifted to be between 0 and 1, with a value of 0 corresponding to identical multigraphs at adjacent learning steps, and a value of 1 meaning the two states of the network do not share a single transition under any input.

### Four stages of learning

Tracking the rate of topological change over learning reveals initial large synaptic changes, which, however, do not change the dynamics in any qualitative way (*d*_dyn_ = 0). Once some meaningful underlying attractors start to emerge, the network undergoes rapid and substantial reorganization of the attractor structure, which brings with it substantial improvements in performance. The attractor structure continues to be adjusted even as behavioral performance saturates finally reaching the stable prototypical cube solution of (25) (Fig. 1c).

These observations can be refined by breaking the learning process into a sequence of *learning stages*. Early in learning (Stage I), *d*_dyn_(*t*) *≈* 0, as the network topology remains the trivial multigraph of a single node, resulting in poor behavioral performance, i.e. ℒ *≫*0. In Stage II, network dynamics begin rapid reorganization (*d*_dyn_(*t*) *≫*0), and behavioral performance improves though is still well above criterion. In Stage III, the network has learned the task adequately (ℒ*≈*0) while the network topology continues to refine (*d*_dyn_(*t*) *≫*0). Finally, in Stage IV, both performance and topology stabilize, as the network finds the universal solution (Fig. 2a). These patterns are robust across a variety of learning configurations, in particular network initialization statistics and learning rate (Fig. S1a).

**Fig. 2.**
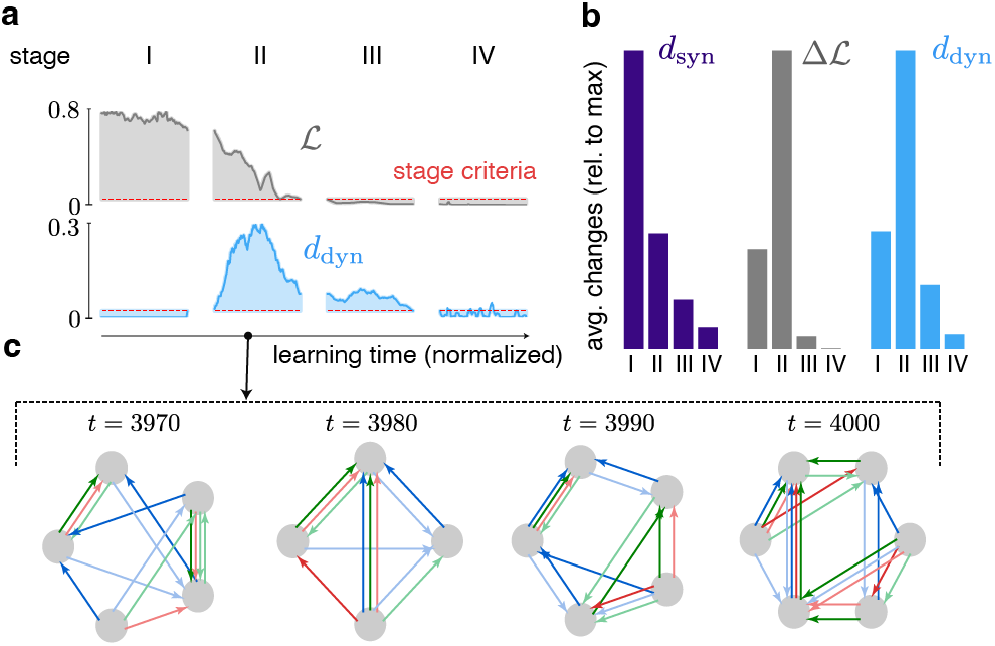
**a)** An example network learning trace, described in terms of the filtered test loss (grey) and the corresponding rate of dynamical change (blue). The criteria used to determine stage assignments shown as red dotted line. **b)** Average rate of gross synaptic change (left, purple) and rate of test loss improvement (right, grey) over time points within stage and over 18 different networks. These 18 networks varied in their random seed for initialization, learning rate used in training (10^*−*2^, 10^*−*3^), and the initial spectral radius of the initialization (0.3, 1, 1.2). **c)** Example multigraphs densely sampled from Stage II, to illustrate what a large rate of dynamical change looks like, featuring both changes in the total number of nodes and the pattern of transitions among them.

The existence of Stage I represents at minimum a theoretical curiosity about how accessible complex dynamics are to randomly initialized networks—large changes must be made in the space of **W** before reaching regions where nontrivial fixed points emerge (26). But beyond that, its contrast with Stage II offers proof of concept for how learning-related changes in synapses can dissociate from behavioral improvements, which are more tightly coupled to changes in the underlying dynamics.

Stage II is crucial for task learning, in two senses: it is the period of largest changes in behavioral performance (Fig. 2b), and the period of largest qualitative changes in network dynamics. Interestingly, this period is not particularly notable from the perspective of synaptic strength changes, which are smaller in magnitude than during Stage I. Nonetheless, these relatively modest changes in **W** drive significant changes in the network functionality. Even on the timescale of one or two bit flips (Δ*t* = 10), these can manifest as changes in the number of nodes or as a complete reorganization in the structure of input-driven transitions (Fig. 2c). This rapid, disorderly evolution of the network dynamical structure (similar to 27) drives a substantial improvement in behavior, suggesting that meta-dynamical analyses offer more direct insight into learning than raw synaptic plasticity.

Stage III is less important for behavioral improvement— which, by definition, has already stabilized to nearly perfect performance—as it is for mechanistic fine-tuning. It has been shown that RNNs of different architectures converge to the same solution, even though the objective function is strictly based on task performance and does not account for the “how” of the solution (25). Nonetheless, learning as driven by behavioral error effectively encourages the network to find not only any dynamics that work, but a particularly elegant (and robust) solution. Stage III is where this universal solution gets shaped, using relatively small weight updates (as scaled by the relatively small size of the errors), before the network reaches stability in Stage IV.

### Universality of learning stages

We know there is one universal solution to this task, but to what extent is the pathway to this solution universal? Given the rapid fluctuations in Stage II, we don’t expect these fine details of the learning trajectory to replicate across networks. But at the macroscopic level, is the coarse pattern of meta-dynamics—a slow start followed by rapid changes and fast behavioral improvement, ending with fine-tuning towards a stable solution—a general phenomenon?

With our dynamical distance, we can compare not only the dynamics of a single network to itself at different time points of learning—as in Fig. 3a, which shows *d*_dyn_ between arbitrary time points of learning for an example network—but also make comparisons between the dynamics of *different* networks as well. The top 3 traces of Fig. 3b show ℒand *d*_dyn_ for each of 3 different networks trained by the same learning rule, Random Feedback Local Online Learning (RFLO, 28), a biological plasticity rule based on integrating local eligibility traces with top-down feedback. They each follow qualitatively similar trajectories, highlighted by a clear Stage II featuring large *d*_dyn_ and an associated drop in ℒ. Fig. 3c shows a cross-temporal comparison between two of these networks, highlighting a shared pattern of changes.

**Fig. 3.**
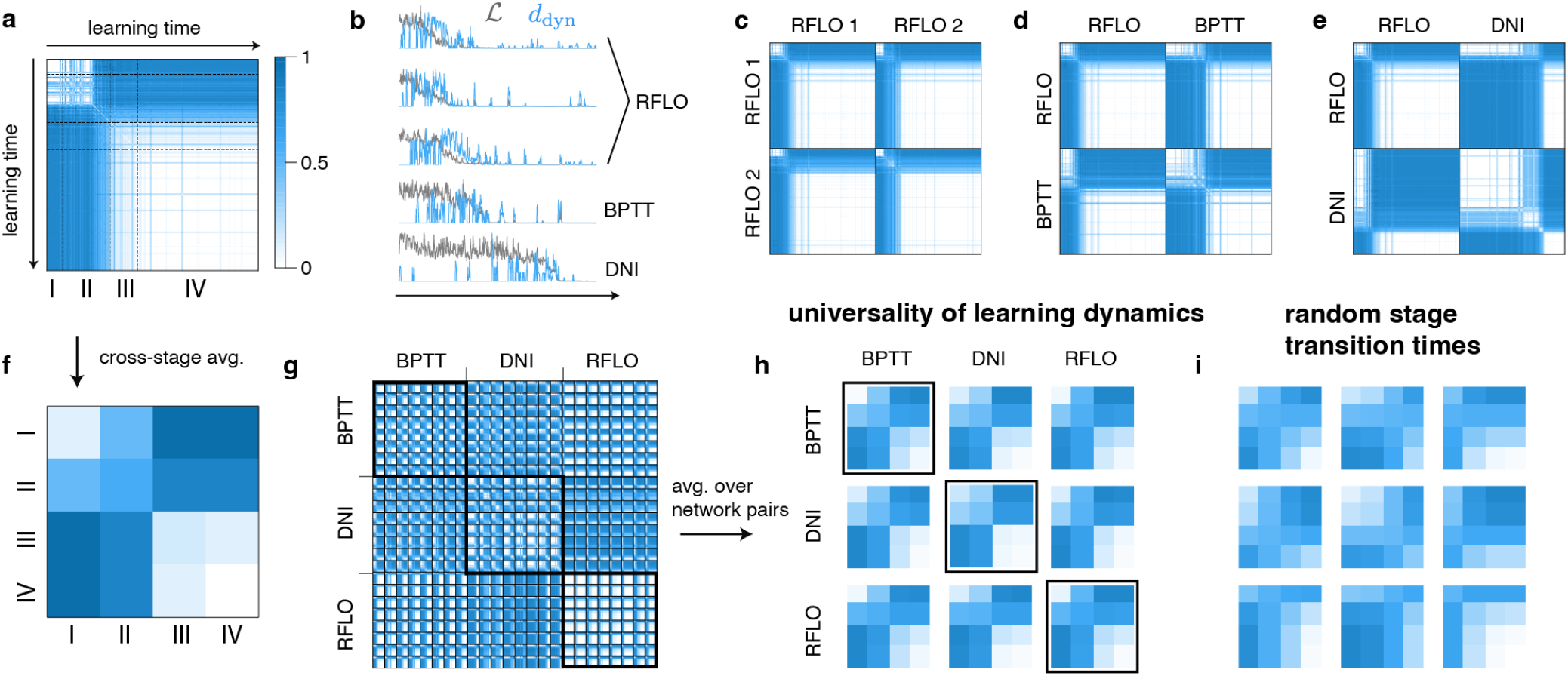
**a)** Topological distance between the state of the network at training time *t*^*′*^ (row) compared with *t* (column). Dashed lines separate the time points into stages. **b)** Example rates of topological change (blue) along with test loss (grey) for 3 example networks trained by RFLo, and 1 for each of BPtt and DNI. In each case, we see bumps in *d*_dyn_ roughly when the test loss decreases, if at different times. **c)** Same as (a) but 2 different networks initialized via different random seeds, each trained by RFLO, compared to one another at all pairs of training time points. **d)** Same as (c) but one network trained by RFLO crossed with a network trained by BPTT. **e)** Same as (d) but RFLO crossed with DNI. **f)** The values of (a) averaged over time points within each pair of stages, showing a coarse stage-by-stage multigraph similarity. **g)** Raw *d*_dyn_ matrix for all 24 networks, generated by 8 random seeds over 3 algorithms. By visual inspection, the block diagonals corresponding to each algorithm are distinguishable, showing differences in the timescale of learning as seen in (b). **h)** Averaging each network block across stages as in (f), then averaging the result across random seeds, we see a common pattern for the meta-dynamics, regardless of algorithm. **i)** Generating random stage transition times, the common structure from (h) is destroyed.

This pattern can be compactly visualized by computing the average dynamical distance ⟨*d*_dyn_ ⟩ among all pairs of time points within bins according to their stage identity (Fig. 3f). Networks in Stage I, typically characterized by the trivial multigraph or other low-node variants, are quite similar to each other and typically quite different from the more sophisticated networks in other stages, as demonstrated by the low value of ⟨*d*_dyn_ ⟩ in the I-I diagonal, but larger off-diagonal values in the row (and column) for Stage I. Stage II multigraphs are highly variable, and they are also quite different from those of other stages, as every entry in the row for II has a large value of ⟨*d*_dyn_ ⟩, including the diagonal. Given that networks in Stage II are undergoing rapid and disorderly changes to their internal dynamics as illustrated in Fig. 2c, we would not expect strong within-nor across-stage similarity. The 2 *×* 2 block diagonal for Stages III and IV has ⟨*d*_dyn_ ⟩ close to 0, showing both that these stages feature little within-stage variability and that they are quite similar to one another: Stage IV networks almost all implement the universal solution, with Stage III networks not far off. Beyond this one example network in Fig. 3f, we see a similar pattern both within and across other networks trained by RFLO (Fig. 3h, third boxed matrix).

But does this hold for all learning processes? We investigate how networks under different learning rules traverse through the space of multigraph structures to arrive at the universal solution. There are many different learning rules for updating synaptic strengths that operate on fundamentally different principles (14). We use three learning rules to train networks on this task and analyze the cross-similarity of their multigraph trajectories over learning. At one extreme, Backpropagation through Time (BPTT, 13) provides an ideal solution for optimizing task performance. But BPTT requires computing complex, high-dimensional learning signals, which make it biologically unrealistic. An efficient approximation to BPTT, the Decoupled Neural Interface approach (DNI, 29) involves learning to compute learning signals that align partially with those of BPTT; it can in principle be implemented under biological constraints, in particular locality of plasticity rules (14). And our third learning rule is the previously introduced RFLO (28), which is intrinsically local. The networks learn the task adequately under all 3 learning rules. However, we can see there are clear differences in the dynamics of learning. The bottom 2 traces of Fig. 3b show behavioral improvement and meta-dynamics on slower timescales than for the networks trained by RFLO. Moreover, Fig. 3d,e show clearly distinct patterns of evolution over the same learning time in direct comparisons of networks trained by different learning rules. Fig. 3g contains a matrix of all cross-network, cross-time dissimilarities in the dynamics, with the large block matrices corresponding to different learning rules (three), the smaller block matrices (eight per algorithm) corresponding to individual networks, and each matrix entry corresponding to a particular time point in training. Although details are hard to parse by eye, the cross-algorithm differences isolated in Fig. 3d,e are also present in the larger matrix’s block structure.

Beneath these cosmetic differences, is there latent similarity in the meta-dynamics for the three learning rules? We repeat the analysis of Fig. 3f by binning time points into stages and averaging *d*_dyn_ within each pair of stages, but we do so for every pair of networks (24 total) featured in Fig. 3g. We then average the result within each learning rule block, generating the average stage-by-stage dissimilarity for networks trained by any pair of learning rules (Fig. 3h). At this level of description, all comparisons show the same learning dynamics as articulated in the single-network example from Fig. 3f. Thus the learning process for the Flip-Flop task is itself also “universal” in this sense.

This universality is not a given *a priori*, nor a subtle circularity inherited in our formalization of the learning stages. The stage transition times are defined strictly by within-network meta-dynamics and behavior, but the matrices in Fig. 3h represent a synthesis of much richer cross-network, inter-temporal comparisons. Our formalization of stages effectively functions as a coarse time-warping of the meta-dynamics to correct for different learning rates. Crucially, it does depend on our stage transition times meaningfully identifying different phases of the learning process for each network. If instead we chose uniformly random stage transition times, the structure is dispersed (Fig. 3i).

There are qualitative elements of the meta-dynamics that distinguish between learning rules, besides the timescale (Fig. 4a). The types of multigraphs generated during Stages II and III are systematically different for RFLO than for the other learning rules. In particular, RFLO networks build up to the 8 required stable nodes for the universal solution must faster and even *overshoot* before pruning the multigraph structure down (Fig. 4b). As RFLO learns the fastest of the 3 approaches, this may be a manifestation of fast and efficient learning in general: the rapid development of an overly complex dynamics, which can be trimmed down to meet the task demands.

**Fig. 4.**
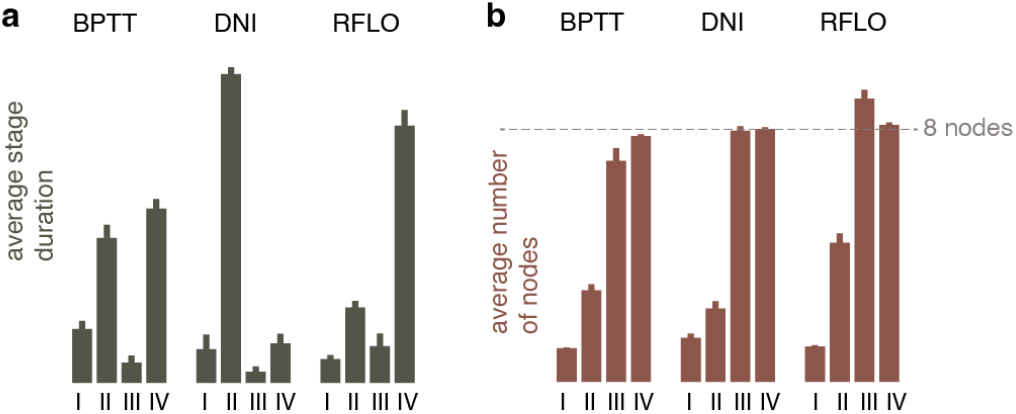
**a)** Average stage duration for all 8 networks trained by each algorithm. Total training time is constant across networks. **b)** Average number of nodes over time points within each stage, averaged over all 8 networks trained by each algorithm. Horizontal dashed line denotes the optimal value of 8 nodes, which training via RFLO exceeds during stage 3.

In summary, networks trained on the Flip-Flop task via different learning rules share a common trajectory towards task mastery, as summarized in 4 stages of learning. Despite idiosyncratic differences among networks in the particular multigraphs they build as intermediate steps, and differences in the rate at which these learning rules achieve success, there is a latent similarity in the pattern of meta-dynamics that leads to learning. A crucial feature of this pattern is the coincidence of multigraph reorganization with the period of steepest improvement in performance, occurring in Stage II. We now turn to other tasks to see if this principle holds more broadly, that meta-dynamics are fundamental for driving behavioral learning.

### Context-dependent evidence integration

The key dynamical features of the Flip-Flop task are fixed points, but these alone do not explain neural mechanisms behind many classic experimental paradigms. For example, line attractors are involved in parametric working memory and evidence integration (6). The emergence of such features complicates the distinction between a topological vs. geometric change in the dynamics. The line attractor is a qualitative feature that is generically useful and distinct from a fixed point, but it also has a spatial extent and orientation, which can grow and rotate continuously.

To study the meta-dynamics of line attractor formation, we train networks on a context-dependent evidence integration task (CDI). Inspired by the paradigm in (10) for context-dependent decision-making, the network is trained to integrate one of two independent, noisy streams of input, depending on which input is cued for that trial (Fig. 5a, CDI). We used both BPTT and RFLO to train networks on this task; DNI is successful on this task to some degree as well, but tends to create degenerate solutions (not shown). In this task, networks develop a line attractor for each context, which is used to integrate the evidence, as seen in noiseless probe trials for different levels of input coherence (Fig. 5b).

**Fig. 5.**
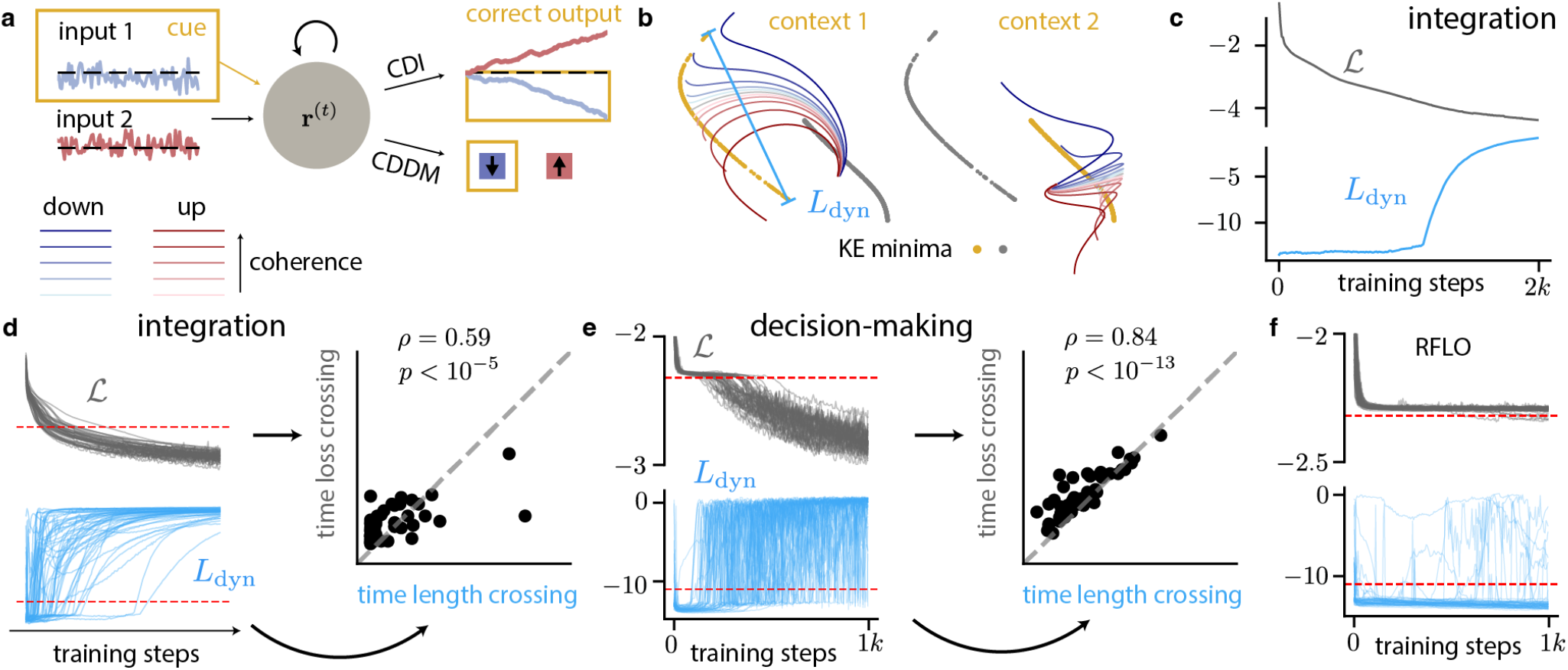
**a)** Schematic of the context-dependent integration and decision-making tasks. Two noisy input streams, a tonic context cue, and a randomly timed report cue (CDDM only) are inputs to the network, from which it must report down/up when prompted by the report cue (CDDM) or the level of net evidence throughout the trial (CDI). **b)** Example trajectories of network activity during noiseless probe trials of CDI, in context 1 (left) and context 2 (right). Dark (light) blue traces represent trials with strong (weak) downward input, and dark (light) red traces represent trials with strong upward input. Yellow dots represent KE minima in the cue context, while grey dots represent KE minima in the uncued context. **c)** Meta-dynamics for an example network trained by BPTT on CDI, featuring the log scale test loss (grey) and the length of the largest node in the system (blue) **d)** Left: Same plot as (c) but for 50 different networks of varying random seed for initialization and training data generation, with thresholds used to separate learning speed and attractor growth (dotted red line). Right: time of threshold crossings for *L*_dyn_ plotted against threshold crossings for *L*. **e)** Same as (d) for CDDM task. **f)** Same as left panels of (d) and (e), for networks trained by RFLO.

How do these line attractors emerge over learning? To assess the meta-dynamics, we use the line attractor *length* as an appropriate metric for mechanism development in this task. That is, we take the set of fixed points found via KE minimization, and within the clusters determined by DBSCAN (see Methods), we measure the spatial extent of each cluster by the maximal distance among pairs of points. The longest length among the clusters gives the dynamical metric *L*_dyn_. We plot it in the same color as *d*_dyn_ from the Flip-Flop task due to its analogous role in measuring critical dynamics-related information—although the metrics are fundamentally different, in measuring geometric vs. topological aspects of the dynamics.^1^ For small values of *L*_dyn_, all nodes are essentially fixed points, with nonzero *L*_dyn_ due to imperfections in the KE minimization process.

Fig. 5c shows an example network’s learning trajectory, with test loss and *L*_dyn_ both plotted on logarithmic scales. The loss decreases gradually over learning, and at some point the line attractor length grows exponentially in size, in a sudden manner suggestive of a phase transition. The relatively dramatic change in *L*_dyn_ contrasts with the gradual improvement in behavior—is there any sense in which the dynamics changes are correlated with behavioral performance, as we saw in Fig. 2b?

Comparing the same curves for 50 example networks, all trained in the same way, but for different realizations of the noise for network initialization and training data generation, we see a distribution of learning speeds (Fig. 5d). We sort these networks by the time at which they cross a loss threshold on one hand, and on the other the time at which *L*_dyn_ grows beyond a threshold, chosen to identify the emergence of nontrivial dynamical systems structure. These event times have a highly significant Spearman rank correlation of *ρ* = 0.59 (*p <* 10^*−*5^), with pairs distributed near the identity line (Fig. 5d). We see similar results for networks trained by RFLO (not shown). This observation suggests that development of network dynamics and behavioral improvement are tightly coupled, recapitulating a key insight from the meta-dynamics of the Flip-Flop task, in a fundamentally different context.

### Context dependent evidence integration with a decision prompt

We analyze the meta-dynamics for a third task, context-dependent decision-making (CDDM), to bridge together the topological and geometric elements of dynamical systems development from the Flip-Flop and CDI tasks. The task is essentially the same as the CDI task, but with an extra input used to cue a binary report of whether the (contextually cued) input is down or up, i.e. net negative or positive. This report cue comes at random times between halfway through the trial and the end of the trial.

Despite the simpler binarized output, this task is more difficult than the integration one, as there are no useful learning signals until the end of the trial, during the report period. The network must learn the long-range temporal dependencies between the correct report and the many pulses of evidence coming earlier in the trial. Moreover, the inputs and outputs themselves only implicitly relate to one another through integration, so the network must figure out the utility of the line attractor representation of evidence on its own, whereas in CDI it is directly “spoon-fed” this computation by the task demands.

Figure 5e shows the learning dynamics of networks trained on the CDDM task by BPPT. Each log test loss curve has an initial sharp decent, which we believe corresponds to calibration of the marginal output statistics. Then each network plateaus for a variable period of time, before eventually breaking through and continuing to improve performance at an exponential rate. Again we sort these networks by the time at which they cross this common test loss threshold, as well as the timing of the line attractor growth. In this case, we see an even stronger and more robust Spearman rank correlation (*ρ* = 0.84, *p <* 10^*−*13^) with these events, and more tightly localized to the identity line.

Our interpretation is that the network must solve long-range temporal credit assignment via BPTT to learn the useful dynamics of evidence integration as a means to an end—even though the task does not explicitly demand a report of net evidence at any point. It is random at what exact time the network discovers this strategy, but once it builds a line attractor, the network is able to quickly accelerate its learning. This interpretation is validated by the null case of RFLO (Fig. 5f), which fails to fully learn the task, likely due to the temporal dependencies exceeding its viable learning horizon. Networks trained by RFLO cannot transcend the same loss threshold from Fig. 5e, and the largest attractor features are almost entirely fixed points with no spatial extent. Although a small number of line attractors incidentally get created in the process by chance, there are significantly fewer than in the population of networks trained by RFLO as compared with BPTT, and the networks are unable to employ them in task-relevant evidence integration.

Overall, the CDI and CDDM meta-dynamics tell a similar story as those in the Flip-Flop task, even amid surface-level differences. In all cases, there is an initial delay in the development of dynamical systems structure needed to solve the task, mirrored by initial stagnation of the loss. Break-throughs in network performance align in time with mean-ingful changes to network dynamics, whether the pattern of input-driven node transitions of the Flip-Flop task or elongation of line attractors in CDI and CDDM. This suggests that these meta-dynamics are causal to learning at behavioral level, and thus a natural target for measurable neural correlates of learning.

## Discussion

Despite exciting experimental efforts in building causal approaches of linking synaptic plasticity to behavioral measures of learning, bridging the gap between the two remains a challenge (30, 31). In parallel, theories for learning algorithms are mushrooming (32–34), with little experimental evidence to disambiguate between them. We desperately need new ideas to approach the question of learning and its neural correlates, in ways that are more experimentally tractable. Here we argue for meta-dynamics as a useful lens onto behaviorally relevant learning. Our approach reveals that attractor dynamics and their evolution over learning turn out to have considerable explanatory power over task performance, particularly in the timing of behavioral improvements with respect to changes in the underlying dynamics. Successful learning rules tend to drive similar meta-dynamics, while failures to learn manifest as the inability to develop the required dynamical systems structure.

Our framework highlights core dynamical systems features that carry the underlying computations needed for behavior, with the hope that this representation may transcend differences between artificial and biological networks. This thinking is strongly influenced by (25), with a few key methodological advances. First, our approach goes beyond characterizing autonomous network dynamics, by describing input-driven transitions among stable fixed points. It is precisely these input-driven transitions that causally determine behavior in the Flip-Flop task, and thus they make the natural target for a meta-dynamics analysis. Second, geometric drift and phase transitions make quantifying changes in attractors structure over learning particularly difficult. We had to develop new methodology for a principled alignment of multigraph nodes across stages of learning. Finally, by virtue of the multigraph abstracting away individualities of a network to summarize the essence of what it does computationally, our fixed point alignment procedure could also be used to compare different networks with potentially vastly different physical realizations.

If asked to reverse-engineer a learning algorithm among an *a priori* specified set of options and with full knowledge of the system’s properties, network activity in the trained model is enough to identify the true underlying learning rule (35–37). Nonetheless, this is arguably not a scenario of practical relevance for systems neuroscience: many unknowns mean we can only look for qualitative rather than quantitative distinctions. We have shown that, at the level of attractor meta-dynamics, the similarities trump most idiosyncratic differences across learning algorithms, as shown in (36) with respect to the end solution. However, we don’t claim that this is always true: there may be other distinctions in the nature of the learning process that *can* influence the meta-dynamics in meaningful ways. The nature of the task is one of those; some have a universal solution, other show substantial across-subject variability in computational strategy (23). Different learning algorithms may differentially bias among such solutions, providing richer opportunities for experimental validation. Just from the endpoints of learning, it is possible to distinguish between a gradient-based rule and a reinforcement-based rule by leveraging a brain-machine interface, where the experimenter controls the mapping between neural dynamics and task outcomes (38). With this additional access to the learning objective, one can develop testable predictions for how neural dynamics should change under one rule versus another. Without such access, disambiguating among learning algorithms may require richer learning paradigms, perhaps involving intermediate training steps or other forms of task shaping (39). In this realm, our meta-dynamics framework provides a test bed for designing richer and more informative paradigms for testing theories of synaptic learning.

Our ability to extract theoretical insights into meta-dynamics relies on an unrealistic level of access to the system; an experimentalist cannot exhaustively search the neural state space to reveal its slow points. **N**onetheless, there are steadily improving statistical tools for identifying latent dynamical features such as attractors using large scale population recordings in behaving animals (40–42). **C**ausal interventions, for example via optogenetic stimulation, allow for direct confirmation s tability o f i dentified dy namical sy stems features (11, 20). **M**ore sophisticated experimental tools, for example holographic stimulation, promise richer and potentially more data-efficient characterizations of circuit dynamics, allowing changes thereof to be observed in chronic neural recordings (43). Thus, a testable prediction of our work is that break-throughs in task performance would be associated with qualitative changes in circuit dynamics in relevant brain regions. More broadly, steady innovation in chronically recording and perturbing neural activity make it likely that analogues of our meta-dynamics-based analysis will soon become possible in experiments.

## Methods

### RNN dynamics

We use vanilla recurrent neural networks (RNN) with defining equations

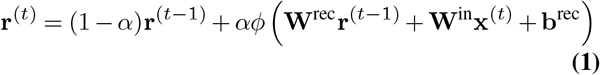

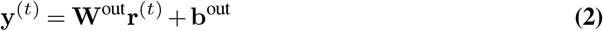

where **r**^(*t*)^ *∈*ℝ^*n*^ is the state of the network, 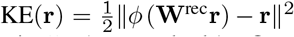in is the task input, and 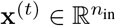 is the network output. The activation function *ϕ* is a fixed, element-wise nonlinearity, and α*∈* (0, 1] parameterizes how time-continuous (α*≈* 0) vs. discrete (α= 1)) the network dynamics are. The trainable parameters **W**^rec^, **W**^in^, **b**^rec^, **W**^out^, **b**^out^ are adjusted by some learning rule (see section on learning rules) to minimize the training error 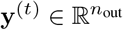, i.e. learn network dynamics where the network output **y**^(*t*)^ matches the desired task output **y**^*\**(*t*)^.

**Table 1.** RNN parameters used throughout the paper.

| Parameter | Symbol | Value |
| --- | --- | --- |
| Inverse time constant (Flip-Flop) | $\alpha$ | 1 |
| Inverse time constant (CDI/CDDM) | $\alpha$ | 0.1 |
| Hidden units | $n$ | 32 |
| Activation function | $\phi$ | tanh |
| Initialization scale | $g$ | 1 |

### Tasks

We used three tasks to evaluate meta-dynamics: the Flip-Flop task, the Context-Dependent Integration task (CDI), and the Context-Dependent Decision-Making task (CDDM). The Flip-Flop task has no trial structure and is simply one continuous stream of inputs **x**^(*t*) ∈^ ℝ^3^. The inputs are i.i.d. sampled at each time step from{*−* 1, 0, 1} with corresponding probabilities of 0.075, 0.85, 0.075. For each of the 3 input dimensions 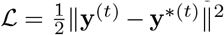, a corresponding output of 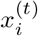 is equal to the most recent non-zero input in that channel. That is, inputs “flip” the output back and forth between 1 and *−*1, which stays constant until the next flip.

For the context-dependent decision-making task (CDDM), we have 5 input dimensions, representing 2 noisy input streams (representing “motion” and “color” information), 2 context cue channels, and 1 probe channel. Each noisy input stream has a mean determined by a coherence value *c ∈*{ ± *×*2^*k*^ 10^*−*3^ : *k* = 5, 6, 7, 8, 9} and a global sensitivity parameter *k* = 0.4. At each time step, the value of the input channel is i.i.d. sampled from 𝒩 (*ck*, 0.1). A trial lasts 100 time steps, the entire duration of which there is a tonic context signal: either 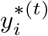 or 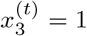 throughout the trial for “motion” and “color” context, respectively, with the other channel set to 0. Finally, the probe input flips from 0 to 1 for a duration of 5 time steps, starting at some random time point between *t* = 50 and the end of the trial. The probe input prompts the network to output a binary decision, for whether the mean input was positive or negative in the appropriate context.

For the context-dependent integration task (CDI), we have an identical setup, except there is no probe channel, and the network must simply report the net evidence at each time point 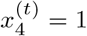 for the cued context stream 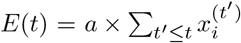, for an integation parameter *a* = 0.025

### Learning setup

We use 3 different learning rules over the course of the paper: Backpropagation through time (BPTT 13), Decoupled Neural Interfaces (DNI, 29, 44), and Random-Feedback Local Online Learning (RFLO, 28). Refer to original papers for algorithm description; here we specify details. For BPTT we use a truncation horizon of *T* = 100 for the CDI and CDDM tasks, matching the trial time. On the Flip-Flop task, which is online in nature and has no natural trial structure, we use the “Efficient BPTT” version of online BPTT from (14) with a truncation horizon of *T* = 6, corresponding roughly to the average period of non-zero inputs. For DNI, we use an affine-linear function of (**a**; **y**^*\**^; 1) to estimate the gradient, with initial weights sampled iid from a gaussian with weight scaling 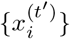. We train the synthetic gradient weights by standard SGD with learning rate *λ*_*sg*_ = 0.003. For RFLO, we used *α* matching the network *α* in Flip-Flop, CDI and CDDM. Crucially, we did *not* project the output error to the hidden state error via random weights, as in the original paper, but rather used the exact projection.

On the Flip-Flop task, we used SGD with momentum parameter *μ* = 0.6, with a learning rate of *λ* = 0.01 for the anecdote in Fig. 2a, and varying between 0.01 and 0.001 for the networks pooled for the analysis of Fig. 2b. For Fig. 3, we used *λ* = 0.003 so that all learning algorithms can learn successfully with identical setup. We also added in L1 and L2 regularization terms with weight 10^*−*4^ on each. For CDI and CDDM, we used Adam with a learning rate of 0.001 and 0.01 respectively, and otherwise standard parameters. For CDI and CDDM, we only used L2 regularization, with weighting of 10^*−*4^.

In the Flip-Flop task, we used online learning (batch size of 1), with a step of learning via the optimizer occurring at every time step of training activity. For the networks in Fig. 3, we trained on 50*k* training time steps per network.

In CDI and CDDM, we used batched learning, with batches of 200 trials per learning step. Learning steps only occur after the entire trial is done. For CDI, we used 2*k* total training steps (with lower learning rate) and 1*k* total training steps for CDDM.

### Fixed point computation

We compute fixed points in the network dynamics by finding minima of the kinetic energy function

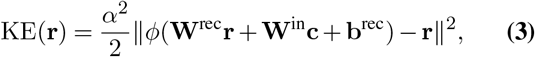

where 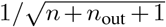 is a tonic context input (as needed in the Context-Dependent Integration task). We minimize via gradient descent over *N*_*GD*_ = 1000 distinct runs from different initial points, resulting in *N*_*GD*_ discovered fixed or approximately fixed points.

The GD initializations are chosen to provide some variability, to find a well distributed set of fixed points in the state space, while also biasing towards parts of state space that are actually used during execution of the task. Thus each initialization point is chosen randomly from network activity during a set of test trials, plus i.i.d. gaussian noise from 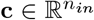.

We begin each run with a learning rate of *λ*. If on 3 consecutive steps, the KE would increase rather than decrease, we would decrease *λ* by a factor of *λ*_*drop*_ and continue. The GD run terminates at the first of 3 stop conditions: a maximum number *N*_*iter*_ of total iterations is reached, *λ* has been reduced *N*_*drop*_ times due to KE increase, or at least *N*_*λ,iter*_ iterations have been reached without reducing *λ*. Finally, fixed points are excluded if their KE is larger than a criterion *KE*_*crit*_. We then take this set of KE minima and identify fixed points and line attractors via the clustering algorithm DBSCAN, with parameter *E*. Unlike *k*-means, DBSCAN keeps line attractors in the same cluster. The centroid of each cluster is used as the representative fixed point for the purpose of mulitgraph computation. We notate the set of all fixed points as 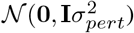.

### Multigraph computation

We empirically assess the stability of these fixed points by running forwards the input-free dynamics from Eq. (1) with initial condition 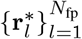 for each *l* and for many random samples of ***η*** *∼ 𝒩* (**0**, *ϵ***I**). At the end of each simulation, we see what other fixed point 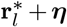 the network state ends up nearest to after *T*_samp_ time steps, and we say 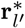 **transitioned** to 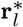 for that random sample ***η***. Sampling ***η*** repeatedly (*N*_samp_ times) gives an estimate of the autonomous transition probability *T*_*ll*_*′* from the *l*th fixed point to each other fixed point *l′*. The subset of points that always transition to themselves are determined to be stable fixed points or **nodes** and retained for the next analysis. Without loss of generality, we index these points as the first *N*_node_ of the initial *N*_fp_ *≥ N*_node_ total fixed points, 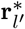.

To compute our multigraph, we compute the above analysis of fixed point transition probabilities, but with two changes:1)as stated above, we include only the fixed point as nodes that are **stable** in the absence of input, and 2) we start each sample trajectory by driving the network with a pulse of input. Starting from all *N*_node_ nodes and probing for 6 different inputs, *±*1 in each of the 3 different input dimensions, generates a set of transition probabilities that define our multigraph: *T*_*pqr*_, the probability of the network initialized near node *p* moving to node *q* under a pulse from input *r ∈ {*1,…, 6*}*.

**Table 2.** Dynamics analysis parameters.

| Parameter | Symbol | Value |
| --- | --- | --- |
| Number of GD runs | $N_{GD}$ | 1000 |
| Initialization variance | $\sigma_{pert}^2$ | 0.5 |
| Initial learning rate | $\lambda$ | 10 |
| LR drop factor | $\lambda_{drop}$ | 0.2 |
| Maximum total iterations | $N_{iter}$ | $10^4$ |
| Maximum LR drops | $N_{drop}$ | 10 |
| Maximum iterations at one LR | $N_{\lambda,iter}$ | $9 \times 10^3$ |
| KE criterion | $KE_{crit}$ | $9 \times 10^{-5}$ |
| DBSCAN distance | $\epsilon$ | 0.5 |
| Number of trajectory samples | $N_{samp}$ | 100 |
| Time steps per sample | $T_{samp}$ | 50 |

### Multigraph comparison

The mathematical problem of comparing two graphs is quite difficult in a vacuum, but it becomes trivial if one can find a principled way to align their nodes. We present a principled way to align the nodes based on their roles in behavior. While it is useful to abstract the nodes away from their locations in network state space and consider the graph as a purely topological entity, we can still retain this information for the purpose of cross-graph node alignment. Although different networks’ latent spaces will not necessarily have some meaningful mapping between them, they do share a common output space since they are trained on the same task.^2^ Thus we project each node 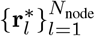 into the output space via its network’s output mapping 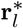. After repeating this projection for the fiducial network with nodes 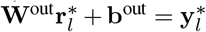 by its own output mapping, we greedily align the pairs of 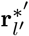 and 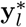 that are closest by Euclidean distance. This produces either a bijective mapping between nodes in the 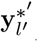 case, or a merely injective mapping in the unequal case.

We then re-shuffle the rows and columns of our network’s transition matrices 𝒯^(*q*)^ to respect the node alignment computed. Then the dissimilarity between these two networks is based simply on the normalized inner product of these transition probability matrices:

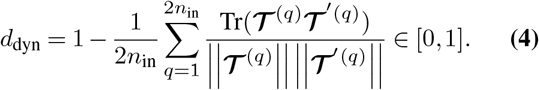

We add in rows and columns of zeros to the matrices with fewer nodes, to match up with unpaired nodes in the larger matrices, if 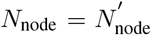. Since these extra dimensions are guaranteed to have 0 absolute alignment, the overall similarity is effectively penalized for differing number of nodes.

### Formal definition of stages

First, irrespective of the order of time points, we do a preliminary classification of each time point as being in Stage I -IV individually. This is based on a temporal averaging of *d*_dyn_ and ℒ_test_ by different rectangular kernels *τ*_*d*_ = 20 and *τ*_*L*_ = 1 (in units of raw time steps), with the former a causal, zero-padded convolution and the latter padded by reflecting over zero. The averaged traces (of same shape as originals) are then compared with thresholds for each. We use *c*_*d*_ = 0.1 and *c*_*L*_ = 0.05 as thresholds for whether *d*_dyn_ ≈ 0 and ℒ_test_ 0 or not, respectively. After a preliminary classification of the time points, we define transition time points from each stage as when *T*_trigger_ = 4 consecutive time points for the next stage appear. Thus to advance from Stage I to II, for example, the filtered *d*_dyn_ trace must exceed *c*_*d*_ for *T*_trigger_ time steps, and then to transition from II to III, the filtered ℒ_test_ trace must dip below *c*_*L*_.

Line attractor analysis

We follow the same KE minimization pipeline as for the Flip-Flop task, up to classification of fixed points via DBSCAN and then separating into stable nodes. We then take, within node class, the maximal pairwise Euclidean distance among its member KE minima, and the (log) maximum of this quantity over all node classes is the value *L*_dyn_ plotted in Fig. 5.

For separating the timing of the loss curves, we used a value of log_10_(ℒ) =*−*3.5 in the CDI task and log_10_(ℒ) = *−*2.32 in the CDDM task. For detecting line attractor growth, we used a threshold value of *L*_dyn_ =*−*11 in both tasks. Due to instabilities in the topology around initialization (likely due to the spectral radius of initialization being near 1), we ignore spurious spikes in *L*_dyn_ if they occur within the first 10 sample networks, out of 200 total (5% of the training steps axis).

## ACKNOWLEDGEMENTS

We’d like to thank Colin Bredenberg for useful discussions. This work was supported by a NYU Dean dissertation fellowship award (OM), a Google faculty award and the National Science Foundation under NSF Award No. 1922658.

## Supplement

**Fig. S1.**
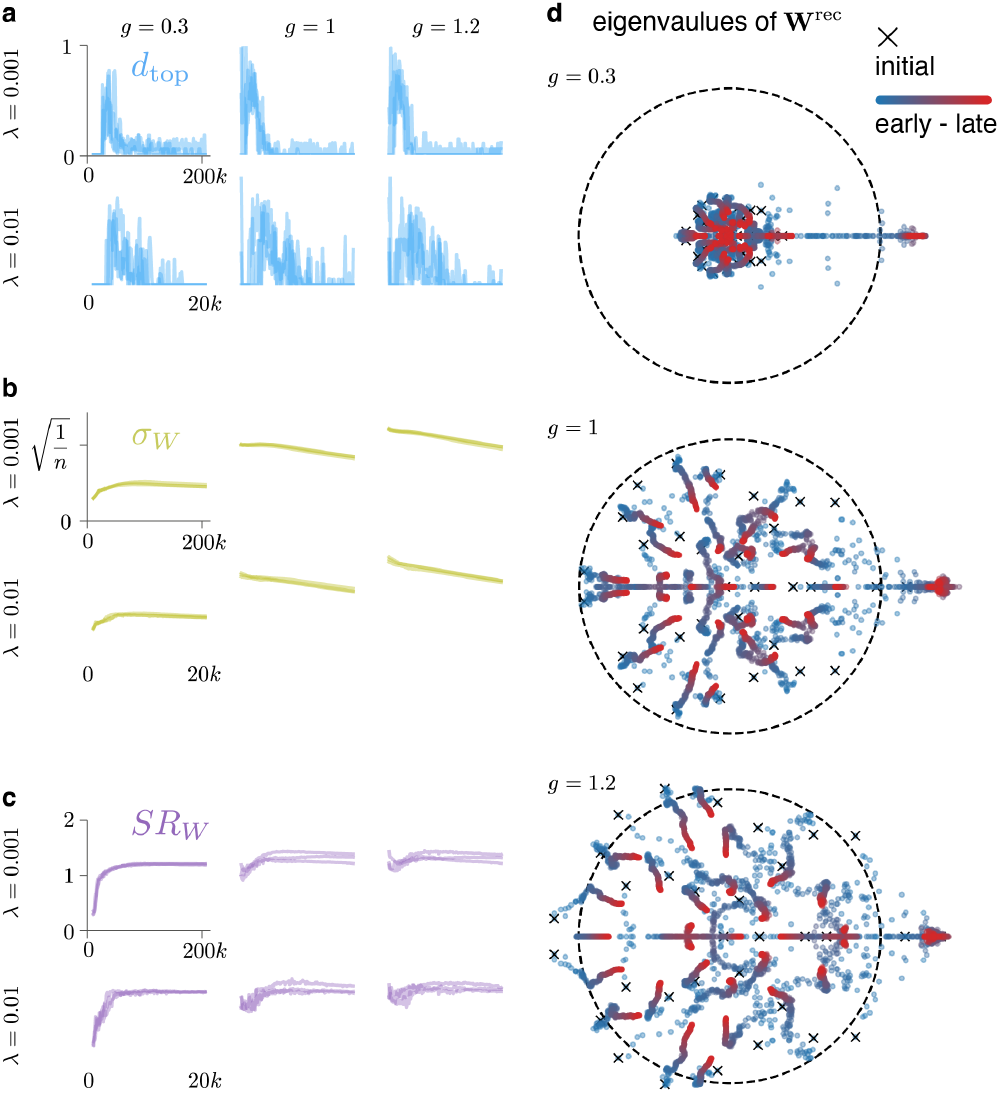
**a)** Many example runs of networks trained on Flip-Flop by RFLO, with different learning rates and different scalings of the network initialization 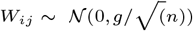. The values of *d*_dyn_ are plotted over learning for 3 example networks per *λ*-*g* configuration. **b)** Same networks, plotting empirical standard deviation of the weights, showing minimal global changes in weight variability. **c)** Same newtorks, plotting the spectral radius, which grows at at least 1 early in training, or was initialized at or above 1 in the case of *g* = 1 and *g* = 1.2. **d)** Plots of the changes in the network spectrum over learning. Early on (blue), the eigenvalues are distributed uniformly over the a circle of radius *g*, and late (red), they mostly shrink towards 0, with the exception of 3 outliers which extend past the unit circle, as in (45).

## Footnotes

1 Moreover, *d*_dyn_ is a distance or rate of change, whereas *L*_dyn_ is a state variable for the network at a given point in learning.

2 They share a common task, but with the caveat that the task is symmetric under permutation of the output indices. Thus the result of this analysis *d*_dyn_ in Eq. Eq. (4) is minimized over all permutations of the output dimensions.

